# The level of specialization of *Phytophthora infestans* to potato and tomato is a biotrophy-related, stable trait

**DOI:** 10.1101/571596

**Authors:** Alexander Kröner, Romain Mabon, Roselyne Corbière, Josselin Montarry, Didier Andrivon

## Abstract

Despite their ability to infect both plant species, natural populations of *Phytophthora infestans*, the pathogen causing late blight on potato and tomato, are usually separated into genetically distinct lineages that are mainly restricted to either host. Laboratory cross-inoculation tests revealed a host-related local adaptation between genotypes, with asymmetric fitness performance between generalist lineages, mainly present on tomato, and specialist lineages confined to potato. To further understand the basis of host-related adaptation in *P. infestans*, we combined experimental evolution and analysis of effectors involved in pathogenicity and cell death modulation. We aimed to check in this way (i) if natural host adaptation of *P. infestans* is reversible during one growing season and (ii) if this process is accompanied by changes in pathogenicity-related gene expression. Two isolates differing substantially by their level of specialization were passaged for nine generations on susceptible potato (cv. Bintje), tomato (cv. Marmande) or alternately on both hosts. Pathogenic fitness and the expression of eight pathogen effectors with known host targets (*AVRblb2, EPIC2B, EPI1, PexRD2, SNE1, PiNPP, INF1 and Pi03192*) and the candidate effector carbonic anhydrase (CA) were quantified before and after experimental evolution on these hosts. Fitness and gene expression varied during the experimental evolution experiment, but independently of the subculturing host. However, the level of host-related specialization of both isolates was stable over time and linked to distinct expression patterns of antagonistic host cell death regulator genes, such as *SNE1* and *PiNPP*. Relations between fitness and effector expression proved to be host- and/or isolate-dependent. Altogether, our results demonstrate host adaptation of *P. infestans* to be a rather stable trait that is not prone to fluctuate by transitory host changes. They further suggest that pathogenicity of *P. infestans* strongly depends on its ability to establish a steady biotrophic interaction with its hosts by regulating effector gene expression.

**Author Summary:** The infamous Irish potato famine pathogen *Phytophthora infestans* causes late blight on potato and tomato, and extensive losses on both crops worldwide. Isolates causing tomato late blight markedly differ in genotype and phenotype from isolates causing potato late blight: under controlled conditions, isolates from tomato perform well on both hosts, while isolates highly pathogenic from potato struggle to produce large lesions on tomato. Mechanisms explaining these differences are unknown, but might provide clues to better understand the fundamental process of host specialization in pathogens. *P. infestans* is known to secrete many effectors, modulating the outcome of the interaction with its hosts. We thus coupled experimental evolution, by subculturing isolates nine times on different hosts, and expression of host cell death regulating effectors to explain pathogenic specialization. We showed that the level of pathogenic specialization depends on the pathogen ability to maintain a biotrophic interaction with its host, and hence to suppress cell death. Host specialization was not altered during serial passages, irrespective of the hosts, although overall pathogenicity increased. These findings show that *P. infestans* is primarily a biotrophic pathogen, feeding on living host tissue, and open ground for new breeding targets for improved resistance to late blight.

## Introduction

Parasitism is the relationship between two species in which one benefits at the expense of the other. Relationships of parasites with plants are diverse, and their classification depends largely on physiological/nutritional and ecological criteria [1–3]. According to the nutritional behavior of the parasite, plant-parasite interactions may be biotrophic or necrotrophic. Biotrophy means that the parasite keeps the host alive and derives energy from living cells (e.g. *Blumeria graminis*) [4], whereas necrotrophy refers to relationships where the parasite kills the host cells to feed on dead matter (e.g. *Botrytis cinerea*) [5, 6]. An initially biotrophic relationship may become necrotrophic (e.g. *Colletotricum graminicola, Sclerotinia sclerotiorum*) [7, 8]. This situation is commonly referred to as hemi-biotrophy, but there is some ambiguity due to divergent use of this term. For example, interactions of the rice blast fungus *Magnaporthe oryzae* are called hemi-biotrophic [9], but there is no single discrete switch from biotrophy to necrotrophy in infected tissue. Biotrophic invasion of host cells is rather sequential and continues in neighboring rice cells while previously parasitized cells die.

Biotrophic hyphae growing ahead of dead tissue have also been reported in the late blight pathogen *Phytophthora infestans* (Mont.) De Bary [10, 11], an oomycete [2] with outstanding scientific and economic importance [12, 13]. Under natural conditions, *P. infestans* depends on living host tissue to complete its life cycle and has low or no saprotrophic survival ability outside its hosts [see 14 and literature cited therein]. However, as tissue damage (necrosis) occurs during later states of colonization, its lifestyle has also been classified as hemi-biotrophic or even necrotrophic. As schematized in Figure 1, the duration of biotrophy varies among *P. infestans* isolates and has been related with the level of pathogenic fitness on tomato in laboratory conditions [15–18]. Biotrophy has also been associated with distinct gene expression patterns on potato [19].

**Fig 1.**
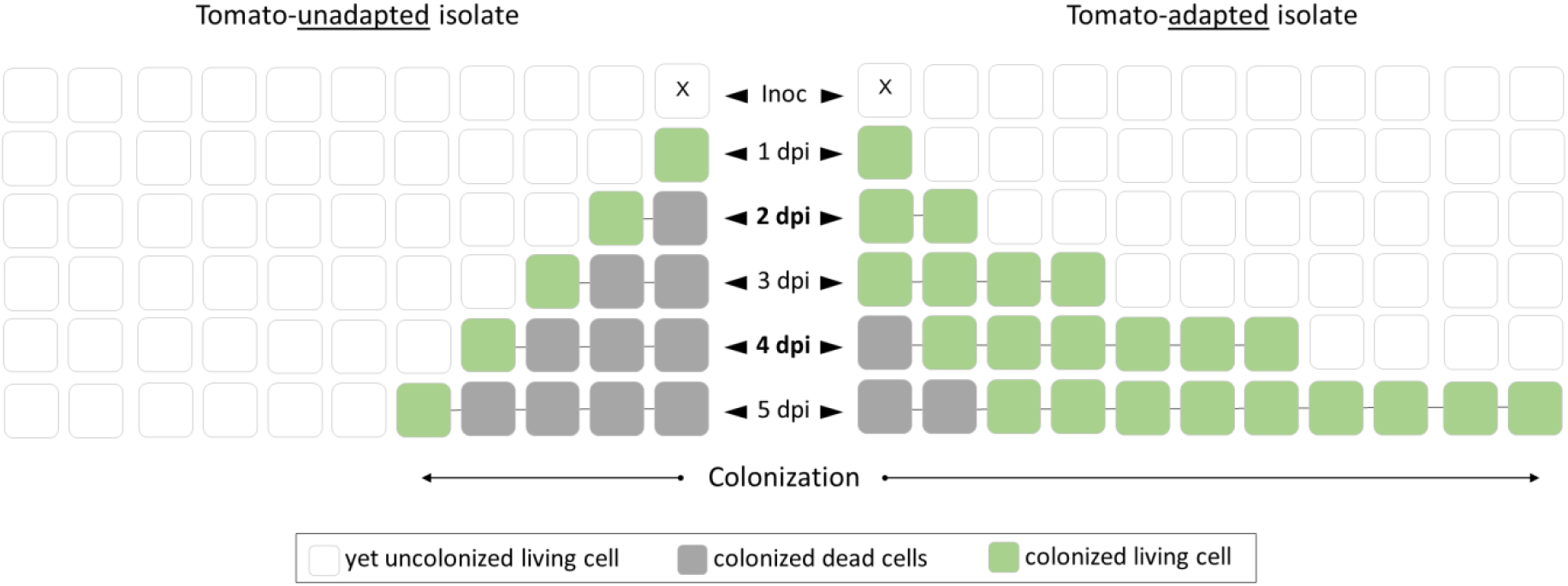
Colonization of tomato leaf tissue by adapted and unadapted *Phytophthora infestans* isolates. This schematic view illustrates spatio-temporal changes of host cell status (uncolonized/colonized, living/dead) in tomato leaf tissue typically observed during colonization by differentially adapted *P. infestans* isolates. Horizontal reading (line-by-line) illustrates the spatial aspect of colonization at one to five dpi respectively. Vertical reading (column-by-column) illustrates the temporal status of a given host cell.

Plant pathogens face in their hosts a two-branched innate immune system [20]: <u>p</u>athogen-associated molecular pattern <u>t</u>riggered <u>i</u>mmunity (PTI) and <u>e</u>ffector <u>t</u>riggered <u>i</u>mmunity (ETI). As outlined by the authors of this seminal review, successful pathogens secrete effectors proteins to suppress PTI and lose/evolve effectors to overcome ETI. These principles also apply to oomycete-plant interactions [21–23]. *P. infestans* is known to employ various effectors during the interaction with its hosts [24, 25] and gene expression polymorphisms among isolates have been reported on potato [19]. Interestingly, these polymorphisms can be associated with the duration of biotrophy [19, figure 7]. *P. infestans* has a repeat-bloated 240 megabases (Mb) genome containing hundreds of fast-evolving RxLR, CRN and other effector-coding genes that are expressed in a coordinated way during its interaction with potato and tomato [24–26]. Secreted into the host apoplast or cytoplasm, they disable host defense components and facilitate colonization: *P. infestans* effectors such as *EPI1* [27], *EPI10* [28], *EPIC1/EPIC2B* [29, 30] and *AVRblb2* [31] are implicated in counter-defense against apoplast-localized host proteases. Other effector genes manipulate host gene expression by downregulating defense related genes [e.g. effector PI03192, 32] or upregulating susceptibility factors [e.g. effector Pi04089, 33]. *P. infestans* also induces/suppresses host cell-death through the coordinated action of cell death antagonistic effectors. As illustrated by Zuluaga et al. [24], biotrophy-related suppressors of cell death such as *ipiO1* [34, 35] and *SNE1* [36] are secreted in the early phases of the interaction, followed by the later secretion of synergistically interacting necrosis-inducing effectors (NIE) such as *PiNPP1.1* and *INF1* [37]. As the timing of expression and biological activity fit trophic phases of the interaction, effectors countering host cell-death may serve *P. infestans* to fine tune the duration of biotrophy by taking control over the host plant cell death machinery [e.g. SNE1/PINPP1.1, 38]. Therefore, *P. infestans* effectors are thought to be involved in host specialization [39]. This view is supported by the fact that *P. infestans* isolates are genetically diverse and may carry different variants of an effector (e.g. ipiO family) [40].

Ecological specialization is a main concept in ecology, but its definitions are highly context-dependent (see [41]). Historically developed as a species attribute (generalist and specialist species), the concept of specialization is now largely extended in the ecological literature to any ecological level (individual, population, species, community). Within-species variation of ecological specialization is indeed widespread [42]: a species considered as ecological generalist may actually be a heterogeneous collection of specialized individuals. Here, we will refer to ecological specialization at the individual level, as the aim of our work is to explore concomitantly ecological and mechanistic aspects of previously documented within-species variation of specialization in *P. infestans* [18, 43, 44]. Considering the important distinction between fundamental and realized specialization [45], it should also be clarified that ecological specialization in the present study refers to Grinnellian specialization assessed in controlled cross-inoculation experiments. Inferred here from direct measures of species performance (i.e. pathogenic fitness) in multiple host environments (i.e. potato, tomato) [46], it analyses the variance in performance across these hosts according to the Levins’ metric [47]. The terms ‘generalist’ and ‘specialist’ refer here to previously identified isolate types (potato isolate, tomato isolate; [15]) showing distinct level of pathogenic specialization.

Overall, what determines the level of host specificity of *P. infestans* is not well understood. Besides the potential effect of cultural practices [48], quantitative differences in pathogenic fitness of isolates may be a particular strong explanatory factor [49–51]. The evolutionary process of pathogenic specialization in *P. infestans* populations on potato and tomato is just as little understood as its underlying determinants. Experimental evolution experiments are thus often used to explore those evolutionary dynamics in real-time. There is evidence, from Mexico and Israel, that tomato isolates could evolve from a local potato population and acquire specialization to tomato [43, 52]. In other words, aggressiveness on tomato would have evolved in isolates already pathogenic on potato. This could explain the lower host specificity of tomato isolates with respect to potato isolates. As documented for various parasite-host systems [53], the observed increase of fitness/virulence/growth on the new host on which the parasite has been passed is often associated with attenuation of these traits on the original host. In the case of highly adaptive pathogens, as little as one [54] or only several passages [55, 56] on the alternative hosts may be sufficient to alter or revert adaptation patterns. *P. infestans* may be considered as a highly adaptive pathogen, as it has been reported to be locally adapted to the host species potato and tomato [15, 18] and generally adapted to particular potato cultivars as well [57–59]. However, to the best of our knowledge, ongoing specific adaptation of *P. infestans* to a particular host cultivar or host species has not yet been established in controlled conditions by experimental evolution experiments. Some evidence arises from an early study of *P. infestans* maintained prior to experiments on chickpea agar [60]. The authors suggest that *P. infestans* could recover initial virulence, but only on its original potato cultivar after serial passages on that specific cultivar.

The present work investigates the lability of specialization in *P. infetans* interactions with potato and tomato using experimental evolution, and explores molecular mecanisms potentially involved in host specialisation. For this purpose, expression of nine effector genes – selected for their established or presumed functions in pathogenicity – were assessed concomittantly to the level of host specialization. Our underlying hypotheses were that i) the original adapation patterns (generalist or potato specialist) could be altered by serial passages through alternate hosts, ii) alterations of pathogenicity towards less host specificity would lower the pathogenicity level on the original host, and iii) each adaptation pattern involves specific combinations or kinetics in the expression of pathogenicity-related effector genes. Our results proved these hypotheses to be wrong, but highlighted the importance of the biotrophic stage in the determination of host specificity in *P. infestans*. More specifically, they showed that the ability for the pathogen to maintain the biotrophic stage for a long enough period is crucial in the sucessful exploitation of the host.

## Results

### Pathogenic fitness of *Phytophthora infestans*

#### *P. infestans* isolates differ in their level of host specialization

Before experimental evolution, both *P. infestans* isolates were equally fit on potato but their level of pathogenic fitness differed markedly on tomato, the isolate collected on tomato (15.TII.02) being fitter than the isolate collected on potato (15.P50.09) (Fig 2A). Each isolate appeared to be fitter on its host of origin than on the other host (Fig 2A). This was most noticeable and statically significant for the potato isolate (*P* = 0.005), but less pronounced and not statistically significant for the tomato isolate (*P* = 0.08). Considering potato and tomato as different ecological niches, the Levin’s measure of standardized niche breath B_A_ (scale 0-1) differed significantly between both isolates (*P* = 0.03), confirming isolate 15.TII.02 (B_A_ = 0.88; se = 0.06) to be more generalist than isolate 15.P50.09 (B_A_ = 0.61; se = 0.04).

**Fig 2.**
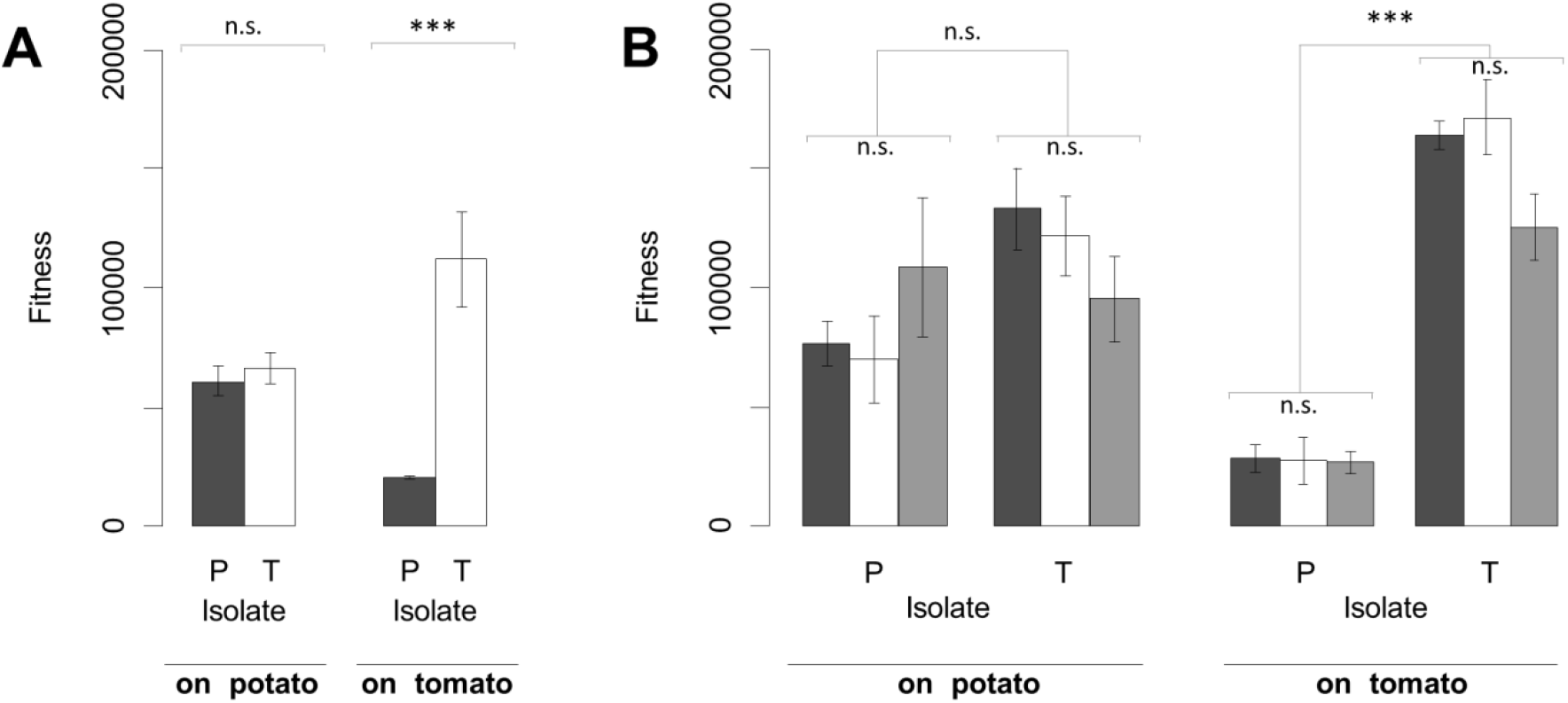
Pathogenic fitness of *Phytophthora infestans* before and after experimental evolution. Two *P. infestans* isolates collected on potato (P-isolate, 15.P50.09) and on tomato (T-isolate, 15.TII.02) were cross-inoculated on detached leaflets of the potato cv. Bintje and the tomato cv. Marmande. Pathogenic fitness at five dpi has been assessed two times: before experimental evolution **(A)** and after experimental evolution **(B)**. Experimental evolution consisted of subculturing the initial inoculum nine times on potato (dark grey bars), tomato (white bars) and alternately on both hosts (light grey bars). Significance of differences was assessed by ANOVA followed by pairwise t-tests (significance level α = 0.05, *** *P* < 0.001, n.s. not significant).

#### Subculturing hosts do not differentially impact pathogenic fitness

Pathogenic fitness of replicate lineages subcultured nine times on either potato, tomato or on alternating hosts did not significantly differ (Fig 2B). This was true for both isolates and whatever the tested host (potato and tomato), showing that subculturing habitats had no differential effect on the pathogenic fitness of *P. infestans* isolates. However, there was a slight general increase of pathogenic fitness during experimental evolution, that was statistically significant only for the tomato isolate.

#### Host specialization is a stable trait (at least over a short-term)

After experimental evolution, the pathogenic fitness of both isolates was still similar on potato and still highly different on tomato (Fig 2B, compare to Fig 2A). As before experimental evolution, the difference in pathogenic fitness of the tomato isolate on potato and on tomato was still less obvious than for the potato isolate. However, it could be shown here to be statistically significant (*P* = 0.02). Despite of this, the initial difference in pathogenic specialization between both isolates was unchanged during experimental evolution, as the evolved tomato isolate remained more generalist (B_A_ = 0.95; se = 0.06) than the evolved potato isolate (BA = 0.62; se = 0.02).

### Effector gene expression

#### State-specific markers and other genes are differentially expressed at 2 and 4 dpi

Transcript abundance differed between 2 and 4 dpi (Fig 3). As expected, transcript abundance of *SNE1* and *PexRD2* genes, involved in biotrophy, was clearly higher at 2 than at 4 dpi (Fig 3A). The same was true for the counter-defense genes *AVRblb2, EPIC2B, EPI1* and *Pi03192*. Transcripts of the stage-specific marker for necrotrophy *PiNPP*, but also of *INF1* and CA, were more abundant at 4 dpi than at 2 dpi (Fig 3B). Increased transcript abundance of these necrotrophy-related genes was associated with the presence of macroscopically visible necrotic lesions at 4 dpi.

**Fig 3.**
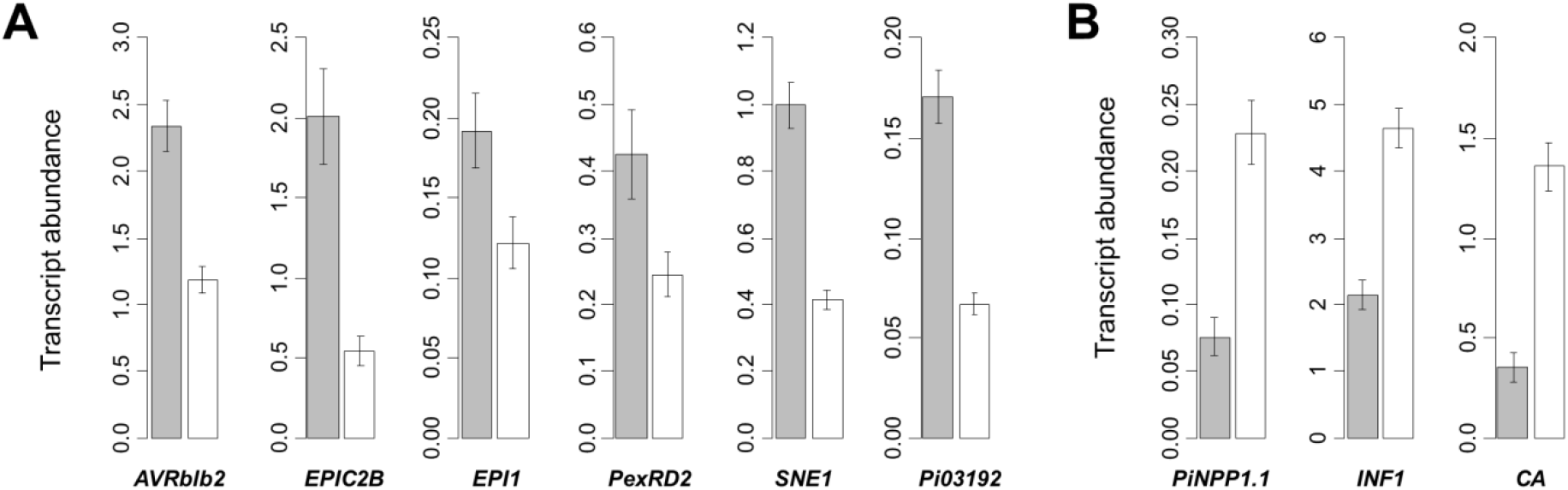
Effector gene expression in *Phytophthora infestans* at two infection stages. Transcript abundance has been assessed at the early biotrophic (2 dpi, grey bars) and at the later transitory/necrotrophic stage of infection (4 dpi, white bars). Some genes were found to be more expressed at 2 dpi **(A)**, but others at 4 dpi **(B)**. The relative timing of gene expression was the same for both isolates (15-P50.09 and 15-TII-02) tested on both hosts (the potato cv. Bintje and the tomato cv. Marmande), before and after nine times subculturing. Results are thus presented as the global mean of these factors (n = 96).

#### Effector genes are differentially expressed between isolates and hosts

Transcript abundance differed significantly between isolates and host plants, revealing distinguishable patterns (Fig 4). Transcript abundance of *EPI1, EPIC2B, CA* and *PiNPP* was significantly higher in the potato isolate than in the tomato isolate, whatever the host (potato and tomato). Transcript abundance of *PiNPP* was also higher in the potato isolate growing on its alternative host, *i.e*. tomato, than on potato (PT > PP), contrasting with the similar transcript abundance found in the tomato isolate grown on each host. Transcript abundance of *PexRD2* was generally higher on potato than on tomato (PP and TP > PT and TT). Interestingly, transcript abundance of *SNE1* was high in the tomato isolate grown on potato compared to other isolate-host combinations (TP > PP, PT and TT). On the opposite, transcript abundance of *INF1* and *AVRblb2* was lower in the potato isolate grown on potato than in other isolate-host combinations (PP < TP, PT and TT). Finally, transcript abundance of *Pi03192* did not significantly differ among all tested isolate-host combinations.

**Fig 4.**
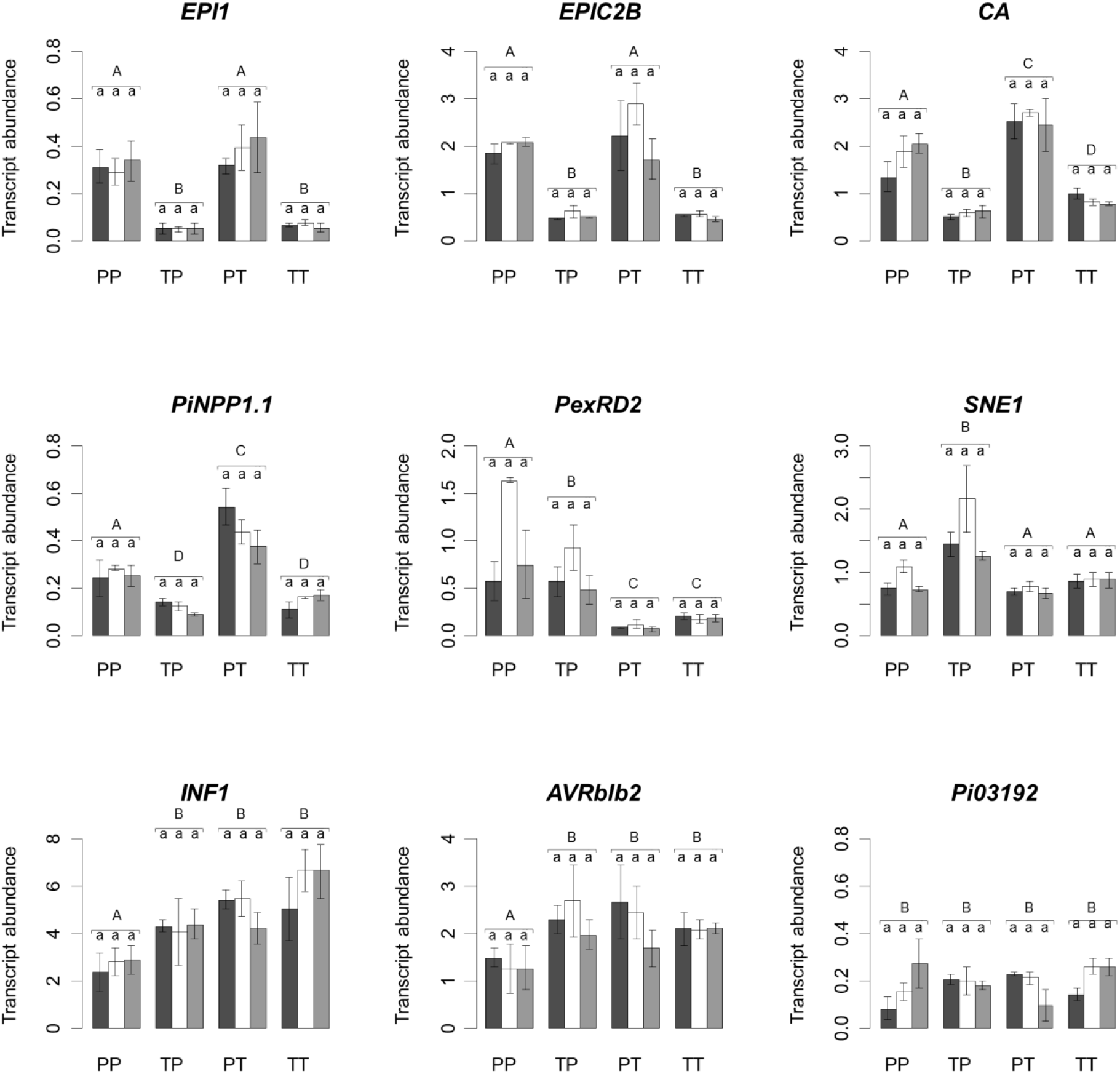
Effector gene expression of *Phytophthora infestans* after experimental evolution. Two *P. infestans* isolates collected on potato (P-isolate 15.P50.09) and on tomato (T-isolate 15.TII.02) have been subcultured nine times on potato (dark grey bars, n=3), on tomato (white bars, n=3) and alternately on both hosts (light grey bars, n=3). Transcript abundance of these experimentally evolved isolates was assessed on the potato cv. Bintje and the tomato cv. Marmande at two dpi *(INF1, CA* and *PiNPP* at four dpi), resulting in the following isolate-host combinations: P-isolate on potato (PP), T-isolate on potato (TP), P-isolate on tomato (PT) and T-isolate on tomato (TT). Different capital letters above bars indicate significant mean differences between isolate-host combinations (pairwise t-test, n=9, significance level α = 0.05). There were no significant differences in transcript abundance among subculturing hosts (see minuscule letters above bars).

#### Gene expression patterns remain stable in experimental evolution on different hosts

Overall, similar levels of transcript abundance were measured in *P. infestans* lineages subcultured on potato, tomato or on alternating hosts (Fig 4). However, it is noteworthy that after subculturing on tomato, transcript abundance of the biotrophy related genes PexRD2 and SNE1 was slightly – but not significantly – increased on potato relative to other subculturing hosts.

### Matching pathogenic fitness and effector gene expression

#### Subculturing on host tissue leads to isolate-specific phenotypic variation

After nine passages on their original host, the pathogenic fitness of the potato isolate was unchanged whereas that of the tomato isolate significantly increased (Fig 5A). This isolate-specific increase of pathogenic fitness was accompanied by a significantly lower transcript abundance of *AVRblb2* and *EPIC2B* but higher transcript abundance of *Pi03192* and *PiNPP* (Fig 5B). No significant variation of these genes was observed in the potato isolate, with unchanged pathogenic fitness. These isolate-specific effects of subculturing were comparable on both tested hosts, and regardless of the host on which they had been subcultured before. Transcript abundance of the other genes tested (*EPI1, SNE1, PexRD2, INF1* and *CA*) was not significantly different before and after subculture (S1 Fig).

**Fig 5.**
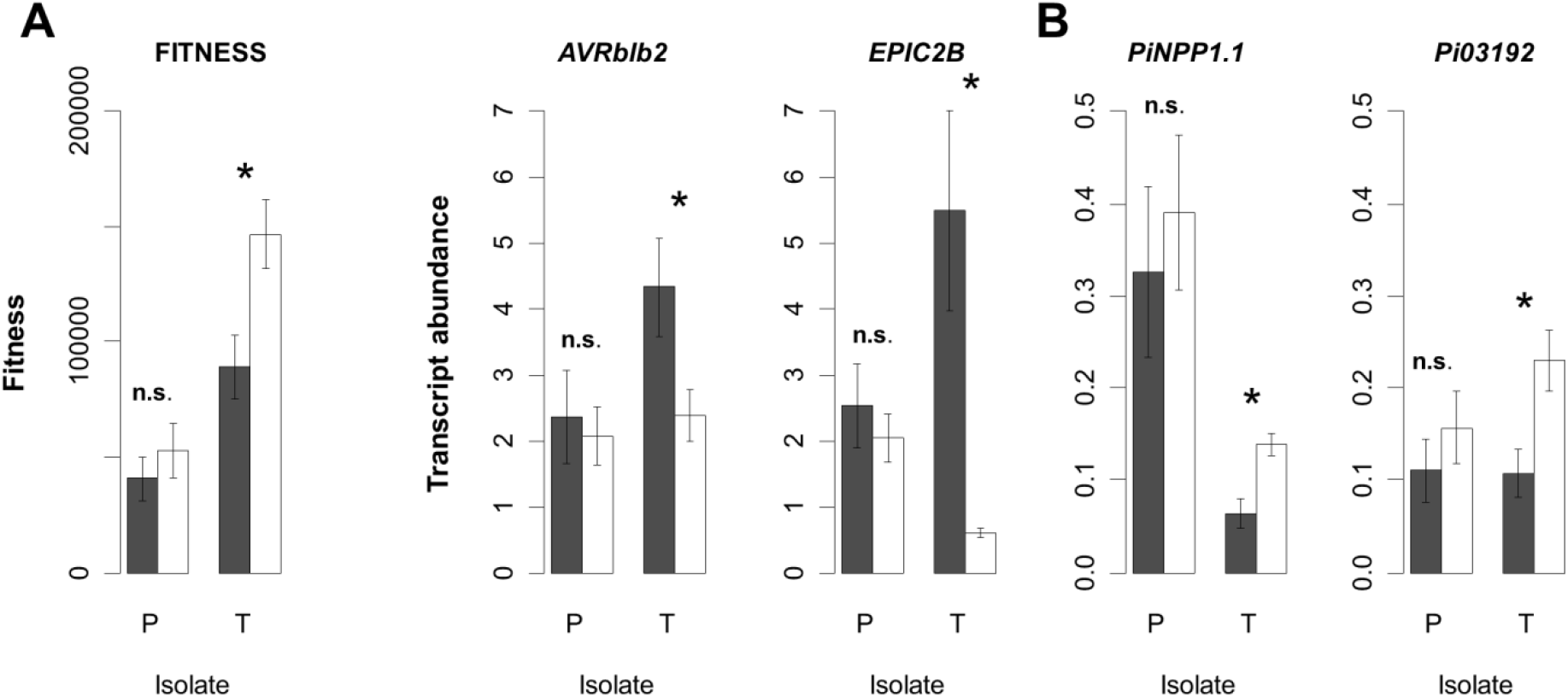
Isolate-specific effects of subculturing on fitness and effector gene expression. Two *P. infestans* isolates collected on potato (P-isolate 15.P50.09) and on tomato (T-isolate 15.TII.02) had been subcultured nine times on their original host: the P-isolate on the potato cv. Bintje and the T-isolate on the tomato cv. Marmande. Fitness **(A)** and effector gene expression **(B)** were assessed before (dark grey bars) and after subculturing (white bars). Fitness was assessed at five dpi and gene expression at two dpi (or at four dpi for *PiNPP*). Both measurements were performed on potato and on tomato. Results were averaged as the effects of subculturing were similar on both tested hosts. Significance of isolate-specific mean differences (t-test comparing before and after subculture, n=6, significance level α = 0.05) is indicated above bars (n.s. = not significant, * *P* < 0.05).

#### Redundancy analysis points to isolate-specific reaction patterns on potato and tomato hosts

Variation of fitness and effector gene expression – generated by the study of two *P. infestans* isolates on two hosts, before and after experimental evolution in different habitats – is summarized jointly by redundancy analysis (RDA, Fig 6). The final RDA model explains 64.11 % of the total variance in the dataset. The first and second constraint component axis represent 56.54 and 19.35 % of the constrained variance (cumulative 75.89 %), respectively. A third constraint component axis (S2 Fig) represents an additional 15.48 % of the constraint variance (cumulative 91.36 %). The first axis separates isolates (Fig 6A; tomato isolate on the left, potato isolate on the right), the second axis displays a separation according to the host on which the isolates were tested (individuals tested on tomato on the top and individuals tested on potato below). The statistical significance of these arrangements – suggesting the phenotype of isolates to be distinct and to depend on the host – was confirmed by a permutation test for RDA (*P* < 0.001). However, the effect of the subculturing host is not significant (*P* = 0.17) and thus not illustrated. Furthermore, significant interaction terms (isolate x host, *P* = 0.003 and isolate x subculture, *P* = 0.002) point to isolate-specific reaction patterns. As obvious from arrangements on the factorial map, the tomato isolate phenotype is distinct before and after subculture (illustrated by open and filled boxes), but the potato isolate phenotype stayed unaltered (isolate x subculture interaction). On the contrary, the model suggests phenotypic variation of isolates on different hosts to be more pronounced for the potato isolate than for the tomato isolate (isolate x host interaction), consistent with the clear host specificity in this isolate.

**Fig 6.**
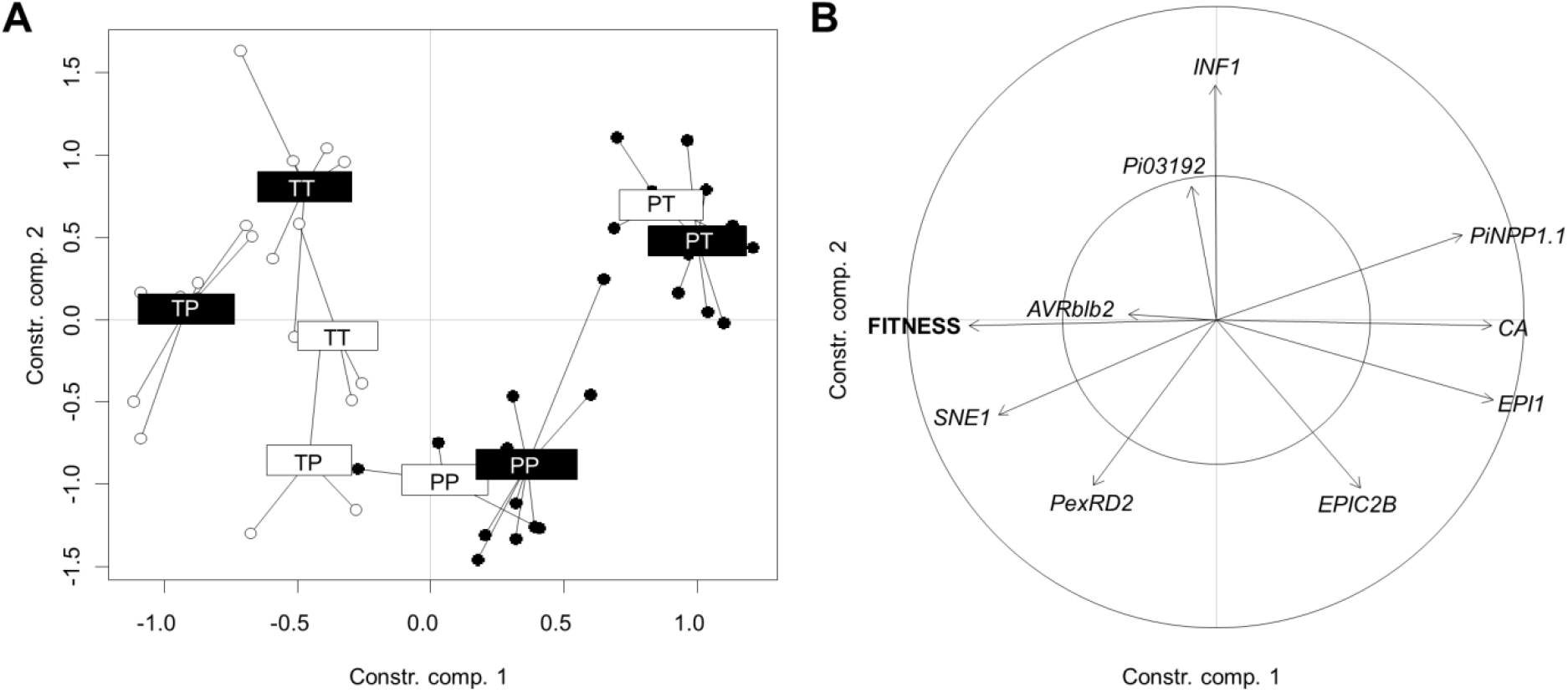
Redundancy analysis (RDA) for transcript abundance and fitness of *Phytophthora infestans*. **(A)** A factorial map separating factor combinations. White boxes represent centroids before experimental evolution and black boxes centroids after experimental evolution. Isolate-host combinations are abbreviated as follows: P-isolate on Potato (PP), T-isolate on Potato (TP), P-isolate on Tomato (PT) and T-isolate on Tomato (TT). Subculturing habitats (on potato, on tomato, alternately on both hosts) are not tagged because model comparison with and without the subculturing host variable revealed no significant difference (*P* = 0.82). **(B)** A loading plot showing correlations among effector genes and fitness.

#### Pathogenic fitness is correlated with effector gene expression

Vector alignments in the loading plot of RDA analysis (Fig 6B) indicate pathogenic fitness to be positively correlated with the expression of *SNE1* and *PexRD2* (grouped left arrows), but negatively correlated with that of *PiNPP, CA, EPI1* and *EPIC2B* (left arrow of fitness opposed to grouped right arrows of effector gene expression). Vectors of fitness and *INF1* are almost orthogonally aligned, indicating no correlation between both variables. The short vector length of *AVRblb2* and *Pi03192* illustrates their low contribution to the RDA model. For verification purpose, RDA analysis was also performed separately on data obtained before and after experimental evolution: vector alignments were consistent in both situations, with the exception of *EPIC2B*. Pairwise correlation tests on unconstrained data (correlations ignoring factor levels) confirmed the statistical significance of all correlative relationships mentioned above (Table 1).

**Table 1:** Pairwise correlations between fitness and effector gene expression of *P. infestans*. Pearson product-moment correlation coefficient r and p-values were calculated on unconstrained data (n = 12, grouped factor combinations). Asterisks indicate significance: p < 0.01 **, p < 001 ***.

| gene | *r* | *p*-value |  |
| --- | --- | --- | --- |
| *EPI1* | -0,65 | > 0,0001 | *** |
| *PiNPP* | -0,59 | > 0,0001 | *** |
| *CA* | -0,58 | > 0,0001 | *** |
| *EPIC2B* | -0,45 | 0,0013 | ** |
| *PexRD2* | 0,40 | 0,0048 | ** |
| *SNE1* | 0,39 | 0,0068 | ** |
| *Pi03192* | 0,15 | 0,2932 |  |
| *AVRblb2* | 0,06 | 0,7018 |  |
| *INF1* | -0,02 | 0,8730 |  |

#### Correlations between fitness and effectors are host- and isolate-specific

Pairwise correlation tests constrained to one host (potato and tomato) or to one isolate (15.P50.09 collected on potato and 15.TII.02 collected on tomato) were performed to explore within-group correlations (Fig 7). Correlations between fitness and effector gene expression were statistically significant on tomato but not on potato (except for *EPIC2B*, Fig 7A). Because a visual inspection of correlative data revealed the presence of split data clouds on tomato (two data groups with different means for each variable), each corresponding to data from one isolate, correlation tests performed separately on data from each isolate revealed isolate-specific correlations between pathogenic fitness and transcript abundance (Fig 7B). *PiNPP* and *INF1* were each positively correlated with pathogenic fitness in isolate 15.TII.02 (collected on tomato), but negatively correlated with fitness in isolate 15.P50.09 (collected on potato). *PexRD2* was not significantly correlated with pathogenic fitness in isolate 15.TII.02, but positively correlated in isolate 15.P50.09.

**Fig 7.**
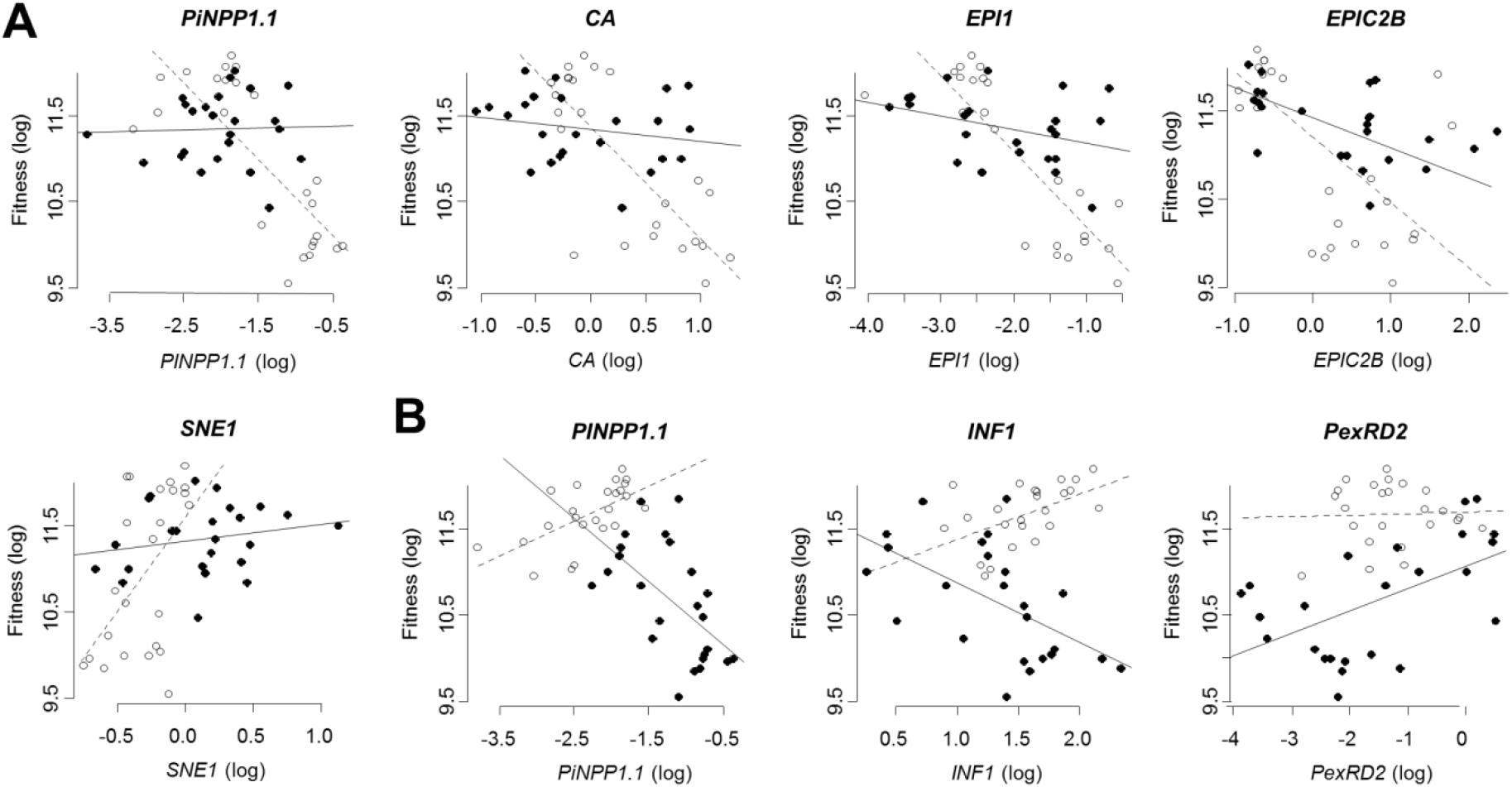
Within-group correlations between fitness and effector genes of *Phytophthora infestans*. **(A)** Data were grouped by tested host. Black circles represent data obtained on potato (cv. Bintje, correlations not significant at α = 0.05, except for *EPIC2B*) and white circles represent data on tomato (cv. Marmande, correlations highly significant). **(B)** Data were grouped by isolate. Black circles represent data obtained for isolate 15.P50.09 (collected from potato, significant negative correlations for *PiNPP* and *INF1* but positive correlation for *PexRD2*) and white circles represent data for isolate 15.TII.02 (collected from tomato, highly positive correlations except for *PexRD2*).

## Discussion

In the present work, quantitative differences in host specialization of *Phytophthora infestans* to potato and tomato were analyzed in terms of stability over serial passages through different hosts and in relation to effector gene expression. The two *P. infestans* isolates included in this study were chosen to display distinct levels of host specialization [15]: the isolate from tomato performed equally well on both hosts (therefore called a generalist), but the isolate from potato, highly pathogenic on its original host, struggled on tomato (therefore called a potato-specialist). As such a pattern has also been consistently reported from other studies [17, 18, 44, 61], the quantitative differences in terms of specialization on potato and on tomato of the isolates studied here may be considered to be representative for *P. infestans* populations on these hosts.

Our work clearly illustrates the absence of ongoing specific adaptation of *P. infestans* to potato and to tomato in controlled conditions: initial differences in pathogenic specialization were maintained throughout the experimental evolution. This may not be intuitive *a priori* for at least two reasons. First, pathogenicity changes during experimental evolution has been documented for various pathosystems, and in many cases the increase of fitness/virulence/growth was specific for the alternative host on which the parasite had been passed and associated with attenuation of these traits on the original host [53]. Depending on the pathosystem, as little as one [54] or only few passages [55, 56] on alternative hosts may be sufficient to obtain such patterns. Second, *P. infestans* is best described as a highly adapted pathogen as it has been reported to be locally adapted to the host species potato and tomato [15, 18] and generally adapted to particular potato cultivars as well [57–59]. There are even evidences that tomato isolates could evolve from a local potato population and acquire specialization to tomato [43, 52]. However, to our knowledge, there is only timid evidence for the possibility of ongoing specific adaptation of *P. infestans* to a particular host cultivar or host species in controlled conditions [60]. Results from our experimental evolution experiment do not contribute on this, but are rather in line with other studies on *P. infestans* stating the absence of ongoing specific adaptation to a particular host cultivar [59, 62] or host species [61, 63] in controlled conditions. We therefore conclude that the levels of quantitative adaptation of *P. infestans* to different host – and therefore the level of host specialization – is a stable trait, at least on the short term and in the conditions of the test. If the here realized number of nine serial passages equals as believed the number of pathogen generations during a growing season in the field, we may speculate that the observed isolate-specific levels of host specialization are not inverted in the field. Most obviously, potato isolates struggling on tomato are not expected to get adapted to tomato during one growing season. We cannot exclude the existence of additional evolutionary forces in natural environments that could accelerate the process of adaptation, but the durable population separation in the field [48, 64] and results from a competitive experiment in field plots [65] are in favor to our view that host specialization is a stable trait. Interestingly, stability of specific adaptation of *P. infestans* to a particular host during the experimental evolution experiment was related to unaltered patterns of concomitantly assessed expression of effector genes.

The comparison of effector gene expression among host-isolate combinations also points to isolate- and host-specific patterns. First, the effectors *EPI1, EPIC2B, CA* and *PiNPP1.1* provided a highly similar profile: their expression was rather similar on both hosts, but differed greatly between both isolates. Second, *PexRD2* and *INF1* were rather similarly expressed in both isolates but their expression level differed between both hosts. These patterns of effector gene expression may reflect isolate and host properties and possibly contributes to variability of pathogenic fitness among host-isolate combinations. Following a validation step on more than two isolates, the observed isolate-specific pattern of *EPI1, EPIC2B, CA* and *PiNPP1.1* could also be used as markers to distinguish both isolate types (isolates from potato, isolates from tomato).

Despite the absence of specific adaptation of *P. infestans* to a particular host, we observed pathogenic fitness after the experimental evolution experiment to be generally increased compared to fitness of the same isolates before experimental evolution. Increasing fitness due to serial passages on a given host has also been observed elsewhere for *P. infestans* on different potato cultivars [66] and from other fungal pathosystems [67, 68]. The general increase of pathogenic fitness could be due to the fact that *P. infestans* was maintained as axenic culture before experiments. It has been established that *P. infestans* loses pathogenicity when maintained on artificial culture medium [60]. The reason for the loss of pathogenicity during axenic culture has not yet been established, but it may be linked to the absence of living host tissue. In fact, in natural conditions, *P. infestans* has a biotrophic lifestyle and no or only limited saprophytic survival ability in the absence of a living host [14, 69]. The increase of pathogenic fitness was only significant for the isolate from tomato. We could thus speculate that the tomato isolate, highly biotrophic on tomato, has lost more pathogenicity than the potato isolate during axenic culture. This could explain the stronger rate of recovery during serial passages on living host tissue. Increased fitness of the isolate at the end of the experimental evolution experiment was related to an altered expression of some of the tested effector genes (*AVRblb2, EPIC2B, PiNPP1.1, Pi03192*). Among these genes, the expression of the protease inhibitor *EPIC2B* was most noticeably altered: it was strongly reduced after experimental evolution in the fitter tomato isolate. Interestingly, no change in effector gene expression was observed for the potato isolate whose fitness was unaltered by experimental evolution. These results show that – even in absence of specific adaptation of *P. infestans* to a particular host – fitness may vary over time and is accompanied by variation of effector gene expression.

Correlation analysis of data for pathogenic fitness and effector gene expression showed biotrophy to be a major clue to explain quantitative differences in specialization of *P. infestans* to potato and tomato. Fitness was found to be positively related with the expression of *SNE1* and *PEXRD2*, two effector genes that have been previously associated with biotrophy in regard to their timely expression and their ability to oppose cell death [36, 70]. As expected by its documented necrosis-triggering activity [37], effector *PiNPP1.1* was also negatively related with fitness in the present study. The antagonistic activity on cell death of *SNE1* and *PiNPP1.1*, supported by our data, has actually been proposed as a potential mechanism for *P. infestans* to control the duration of biotrophy [38, illustrated model, 71-76]. Comparing gene expression in the two isolates included in this study on their respective original and alternative hosts also suggests a similar antagonistic interaction between *PiNPP1.1* and *PexRD2*. Consistent with this view, the potato-specialized isolate coordinately increased expression of *PiNPP1.1* and reduced expression of *PexRD2* on tomato, relative to the expression of both genes on its original host. The resulting stronger host cell death and shorter biotrophic period were associated with a drastically lower pathogenic fitness. This relationship between reduced biotrophy and reduced pathogenic fitness is consistent with the view that *P. infestans* is a biotrophic pathogen that requires living cells to feed on.

Why would potato-specialized isolates of *P. infestans* trigger early host cell death on their alternative tomato host by increased secretion of apoptosis-related effectors (i.e. *PiNPP1.1)* if this is negatively related with fitness? A biological explanation refers to the “increasing plant defense” theory [reviewed by 7], which claims that (hemi-)biotrophic pathogens eventually kill the host cells they infected to protect themselves as a last resort against steadily increasing host defenses. Particularly strong evidence for this theory is provided by work on the fungal pathogens *Magnaporthe oryzae* [77] and *Colletotrichum graminicola* [78]: the authors found that the duration of the biotrophic stage fits the speed of increasing host defense. Our results, as well as literature reports on *P. infestans* [37, 79], suggest that highly expressed host defenses also force this otherwise biotrophic pathogen to actively trigger host cell death (Fig. 8). Literature reports on steadily increasing host defense during *P. infestans* infectious process [80, 81], and their more or less successful effector-mediated suppression [72, 82] strengthen this view.

**Fig 8.**
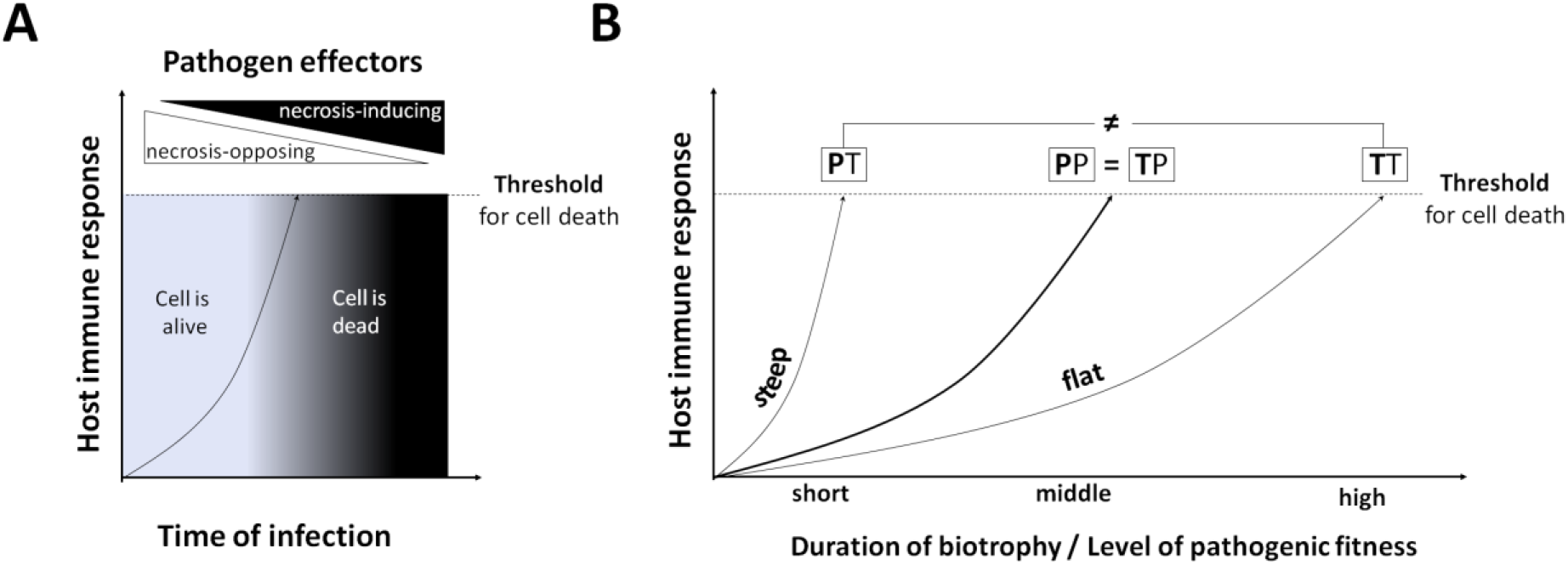
Host immune responses could limit the duration of biotrophy in *Phytophthora infestans* interactions. **(A)** Host immune responses force *P. infestans* to kill its host. It is well established that *P. infestans* secretes effectors into the parasitized host cell adnd that secretion is timely regulated. Biotrophy related effectors are secreted in the early state of the interaction to maintain the host cell alive. With progressing time of infection, secretion of pro-life effectors decreases while necrotrophy related effectors with cell-death inducing activity are increasingly secreted. As the parasitized cells die concomitantly to this switch, a tightly controlled and timely secretion of necrosis-opposing and necrosis-favoring effectors is regarded as a mean of *P. infestans* to regulate the duration of biotrophy. However, the reasons why *P. infestans* – an almost obligate parasite – actively kills host cells is not understood and may even appear counterintuitive. The here advocated explanatory approach refers to the “increasing host defense” theory (see explanations in the text) suggesting biotrophic pathogens to actively kill the host cell as a last resort against an intolerable level (threshold) of host defenses. Literature reports about steadily increasing host defenses in *P. infestans* interactions in conjunction with the here and previously observed timely regulated expression of prolife/pro-death effector genes (e.g. SNE1/PiNPP1.1) support this view. **(B)** Duration of biotrophy in *P. infestans* interactions is positively related with host adaptation. It has been proposed that timing and intensity of host immune responses impact on the outcome of interactions between *P. infestans* and its hosts. Results from a range of studies show that immune responses have to be induced timely and strongly to be effective against *P. infestans*. Considering these results and with regard to the theory that strong host immune responses force *P. infestans* to rapidly kill its host (see A), we would expect duration of biotrophy to be positively related with the level of host adaptation. Consistent to this prediction, similar pathogenic fitness of both tested isolates on potato matched macroscopically assessments of lesions. Furthermore, higher fitness of the tomato isolate on tomato in respect to the potato isolate matched a visibly more biotrophic interaction. Isolate-host combinations are abbreviated as follows: P-isolate on Potato (PP), T-isolate on Potato (TP), P-isolate on Tomato (PT) and T-isolate on Tomato (TT).

In addition to the cell death antagonistic effectors *PiNPP1.1 - SNE1/PexRD2*, we also studied the expression of the counter-defense protease inhibitors *EPI1* and *EPIC2B* [27, 30], that were both found to be negatively related to pathogenic fitness of *P. infestans* on tomato. This relation may be explained through the programmed cell death in hypersensitive reactions (HR PCD), a pathogen-triggered host response to infection that is associated with strong immune responses [83]. The direct implication of *EPI1* and *EPIC2B* in host cell death has not yet been conclusively established, but we argue – based on evidence from literature – that *EPI1* and *EPIC2B* protease inhibitor activity of these effectors could modulate *P. infestans* interactions by blocking immunity related programmed cell death in the host (Fig. 9 and S1 Appendix). This hypothesis is based on two main facts: (i) *EPI1* and *EPIC2B* jointly inhibit protease activity in the tomato apoplast [30] and (ii) at least one of these inhibited proteases *(PIP1)* is required for immunity-related PCD in tomato [84]. We thus hypothesize that *P. infestans* could address PCD during the HR by secreting proteases inhibitors that target cell death-related host proteases. On that condition, expression of proteases inhibitors by *P. infestans* may reflect the intensity of host immune signaling for HR PCD encountered in the tested host-isolate combinations. There is indeed literature evidence that *P. infestans* protease inhibitors are specifically up-regulated in host tissue compared to culture medium [25], and are expressed concomitantly to their respective host targets [27, 30]. We can thus expect a strong host immune signaling for HR PCD in situations where *EPI1* and *EPIC2B* expression is increased, and only a low host immune signaling for HR PCD in situations with low expression of these protease inhibitors. The negative correlation with fitness may be explained by the continuous increase of other immune responses lowering pathogenic fitness. It is now well established that HR is not limited to PCD, but also involves the strong accumulation of various defense compounds and the rapid expression of defense-related genes [85, 86–88]. This apparently also applies to *P. infestans*, where HR PCD is ubiquitous [10, 11, 71, 72, 89] and positively related to timing and/or intensity of defense [71–76]. We therefore speculate that the short duration of biotrophy in the potato isolate-tomato interaction results from a strong host immune response. In the absence of experimental evidence, the severe and fast cell death observed may be either plant-controlled HR PCD and/or pathogen-controlled cell death in response to overwhelming defenses (increasing defense theory, Fig. 8). In both cases, necrosis would be associated with low pathogenic fitness of *P. infestans*.

**Fig 9.**
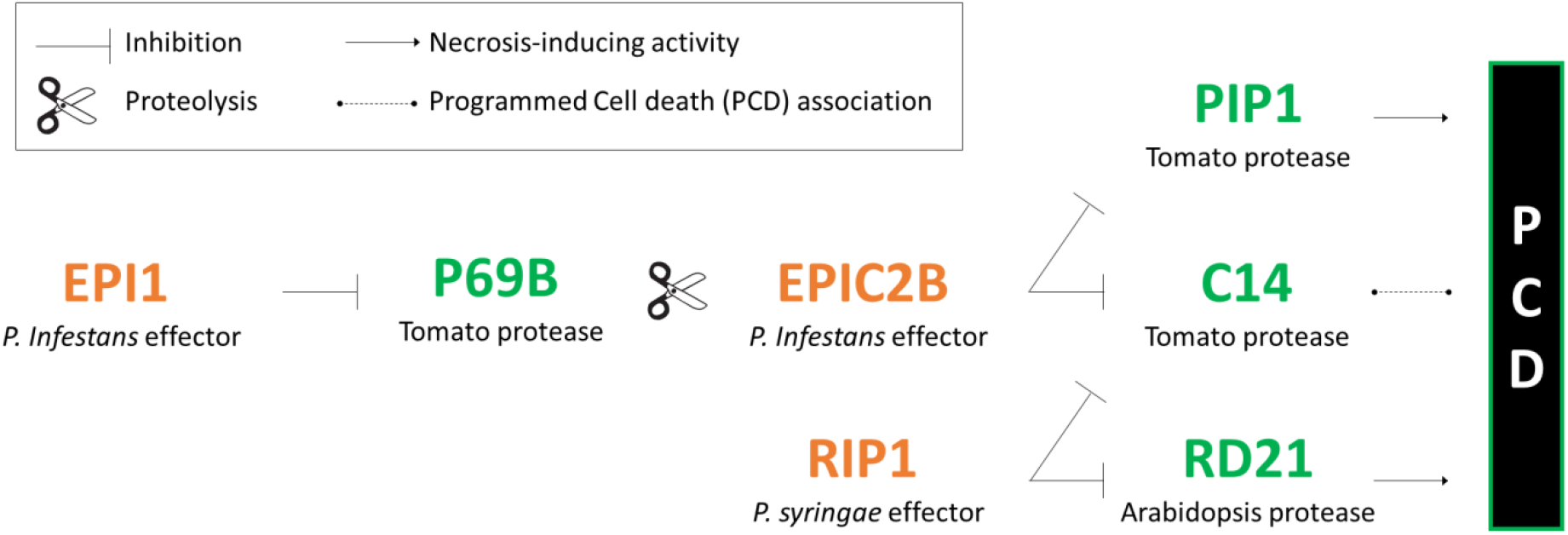
Protease inhibitors EPI1 and EPIC2B could block immunity-related programmed cell death. The *P. infestans* effector EPI1 prevents degradation of EPIC2B by inhibiting the tomato protease P69B. The thereby guarded EPIC2B inhibits in turn the tomato protease C14 that has been reported to be associated with stress-induced programmed cell death (PCD). EPIC2B also inhibits the tomato protease PIP1 that has been shown to be required for HCD. Tomato C14 is furthermore inhibited by the *Pseudomonas syringae* effector RIP1 that also inhibits in Arabidopsis the pro-death protease RD21. Altogether, these literature findings show that *P. infestans* could take control about the tomato host cell death machinery by secreting proteases inhibitors targeting cell death-related host proteases (e.g. PIP1, C14).

To conclude, the set of results reported here clearly shows that the level of pathogenic specialization of *P. infestans* to potato and tomato is a biotrophy-related trait, and that is unaltered by the different tested conditions of our experimental evolution. Our data strongly support the idea that pathogenic fitness of *P. infestans* is positively related with the duration of biotrophy, itself finely regulated by the balance in the expression of antagonistic necrosis-inducing (PiNPP) and necrosis–opposing (SNE1, PexRD2) effectors, and that this balance itself depends on the time required for host defenses to reach a threshold level. Future research may address this hypothesis, by studying inducible host defenses in response to a range of *P. infestans* isolates collected from different hosts. As they stand, these conclusions and hypotheses however carry wide-ranging consequences for the biology and management of late blight, and possibly of similar pathogens (downy mildews in the first place). Indeed, they show that *P. infestans*, often called a hemibiotroph [15–18], should rather be seen as an ‘impecfect biotroph’, since its basic infectious process and trophic mode is biotrophic, but eventually fails to keep its host alive like ‘perfect biotrophs’ (rust or powdery mildews, for instance) do for extended periods of time. The negative association between host damage and parasitic fitness, clearly evident from our data, tend to revert the commonly held equation that increased aggressiveness (*i.e*. faster and more extensive necrosis of host tissue) equates with greater fitness and invasion potential in *P. infestans* ([see e.g. 19]. This, together with the stability of adaptation patterns over serial passages though different hosts, open new and important ways to better control late blight through host genetic resistance.

## Material and Methods

### Plant material

The potato cultivar Bintje and the tomato cultivar Marmande – both highly susceptible to late blight – were grown in a glasshouse as described by Kröner et al. [15]. Fully developed young leaflets were collected from seven week-old plants for experiments.

### *Phytophthora infestans* isolates

The *P. infestans* isolates 15.P50.09 (from potato, EU_13_A2_86 clonal lineage) and 15.TII.09 (from tomato, EU_23_A1 clonal lineage) were collected in 2015 in France from naturally infected potato and tomato crops. They were selected for this study on the basis of previously available genotypic and phenotypic data [15]: they belong to distinct clonal lineages that are – at least in France – typically separated by the host (potato/tomato) and provide the usual pattern of pathogenic fitness *in vitro* on these hosts (tomato isolates perform overall well, but potato isolates struggle on tomato). During six months, both isolates were maintained by serial transfers on pea broth agar medium in darkness at 15°C [90].

### Experimental evolution experiment design

The experimental evolution experiment (S3 Fig) consisted in subculturing *P. infestans* isolates on detached leaflets: either on the original host (from which the isolate had been collected), on the alternative host (e.g. on tomato leaflets if the isolate had been collected on potato), or on alternating hosts (in turns on potato and on tomato). The same inoculum source was used to initiate these conditions for experimental evolution. For each condition, three replicated lineages were subcultured nine times at intervals of seven days. Inoculum (3.10^4^ sporangia.mL^−1^) was prepared in sterile water from sporangia produced on previously infected leaflets. New leaflets were inoculated on the abaxial side with four drops (each 20 μL) of this suspension. A detailed description of inoculum preparation is available in Kröner et al. [15]. During the experimental evolution experiment, temperature was maintained at 18/15°C (day/night) and the incubation chamber was illumined to obtain 16 h day length. Fitness and effector gene expression of *P. infestans* were assessed simultaneously, before and after experimental evolution.

### Estimation of pathogenic fitness

A single fitness estimate, representing the mean fitness of *P. infestans* from three lineages (biological replicates), was calculated as proposed by Montarry et al. [91]. Relevant life-history traits of *P. infestans* (latency, lesion size and sporangia production) were measured on potato and tomato leaflets. Experimental conditions equal those encountered during the experimental evolution experiment, with the exception that only one drop of inoculum (instead of four) was placed next to the center of a detached leaflet. To determine latency period, six replicate leaflets per replicated lineage (6 leaflets x 3 lineages = 18 leaflets in total) were inspected for sporangia formation at 24 h intervals, beginning two days post inoculation (dpi). At five dpi, lesion size and sporangia production were assessed on the same leaflets, as described by Montarry et al. [90]. The two constants of the fitness model were fixed as follows: the underlying leaflet size of potato and tomato (X) was determined experimentally to 20.3 cm^2^, and the assumed time available to exploit the leaf (1/μ) was set to five days. Fitness data (S1 Table) was used to calculate the Levin’s measure of standardized niche breath B_A_ on a scale from zero to one [15, 92]. For these calculations, potato and tomato were considered as possible hosts of *P. infestans*. A low niche breath points to a high level of specialization on one of these hosts, and *vice versa*.

### Effector gene expression

#### Sample preparation

Effector gene expression was assessed at the same time as fitness in order to assure identical conditions: same batch of leaflets, sporangia suspension and inoculation conditions. Leaf discs (22 mm in diameter, centered on the site of inoculation) were sampled at 2 dpi and 4 dpi.

For each replicated lineage (three biological replicates), six leaf discs (one leaf disc per infected leaflet) were grouped, shock frozen in liquid nitrogen and lyophilized. The sample preparation system FastPrep-24 (MPbio) was used to obtain a homogenous powder. This was achieved by three successive 30-second runs at speed 4. The homogenous powder was aliquoted to obtain two 5 mg samples (two technical replicates of crushed tissue) that were stored at – 80 °C prior to RNA extraction.

#### RNA extraction, quality control and cDNA synthesis

For each replicated lineage (three biological replicates), total RNA was extracted separately from two 5 mg crushed-tissue samples by using the SV Total RNA Isolation System (Promega), according to the manufacturer’s instructions. The suitability of this protocol for high quality RNA extraction – in the present experimental conditions – was initially confirmed by calculating the RNA Integrity Number (RIN) from random samples by using the 2100 Bioanalyzer (Agilent Technologies). During routine operation, total RNA was quantified by the NanoDrop 1000 spectrophotometer (Thermo Scientific) and RNA quality was confirmed by agarose gel electrophoresis. Synthesis of cDNA was performed with 1 μg of total RNA by using the GoScriptTM Reverse Transcription System (Promega), according to the manufacturer’s recommendations. The absence of genomic DNA contamination was confirmed by amplification of an intron-spanning region of the elongation factor 1 alpha (EF-1α, GenBank: DQ284495), as described by Rosati et al. [93]. The cDNA samples were stored at −80 °C.

#### qPCR primers, amplification conditions and calculations

qPCR primers for eight *P. infestans* effector genes *(AVRblb2, EPIC2B, EPI1, PexRD2, SNE1, PiNPP1.1, INF1, Pi03192*), the candidate effector gene coding Carbonic Anhydrase (CA) and three reference genes (*ACTA, BTUB, EF2A*) have been designed and optimized for this study (Table S2). Specificity of these primers was checked by sequencing amplicons in both directions. Real-Time qPCR was performed twice on each cDNA sample (two technical replicates of analysis) by using Lightcycler 480 SYBR Green I Master (ROCHE) chemistry in combination with the Lightcycler 480 II system (ROCHE). PCR reaction volumes of 10 μL contained: 1.5 μL PCR-grade H_2_O, 0.5 μL of the forward and 0.5 μL of the reverse primers at a concentration of 10 μM respectively, 5 μL Master Mix (2X conc.) and 2.5 μL of cDNA template. Amplifications were performed in 384-well plates under the following cycling conditions: 15 min at 95°C; 39 cycles of 15 s at 95°C, 30 s at 62°C (but 64°C for *EPI1, ACTA* and *BTUB*) and 30 s at 72 °C. Melting curve analysis was performed from 62°C to 96°C. Transcript abundance of effector genes was calculated by using the relative quantification method described by Pfaffl [94]. To take into account for possible variations of PCR efficiency, separate standard curves from template dilution series were included for each PCR run x primer set x isolate combination (e.g. PCR run 1 x *AVRblb2* primer x isolate 15.P50.09). To calculate transcript abundance, Ct (threshold cycle) values were first transformed to linear scale expression quantities. Expression of the target gene was then normalized in respect to the geometric mean of the three reference genes. These normalized transcript abundance data (S3 Table) are expressed in this article as the means from three replicated lineages of *P. infestans* (three biological replicates).

### Statistical analyses

Statistical analyses were performed using the statistical software R version 3.1.1 [95]. The means of fitness and transcript abundance of effector coding genes were calculated from three replicated lineages of *P. infestans* (three biological replicates), after having averaged – for each replicate lineage – results from technical replicates (for fitness: life-history traits measured on six leaflets; for transcript abundance: two crushed-tissue samples from six leaflets x two PCR analysis).

Null hypotheses were rejected if *P* < 0.05. Data were transformed to logarithms or square roots when necessary before performing analysis of variance (ANOVA), analysis of covariance (ANCOVA), multiple comparisons of means with adjusted P-values (Student’s t-tests), tests for association between paired samples (Pearson’s product moment correlation coefficient) and Redundancy Analysis (RDA, data prepared for multivariate analysis is available in S4 Table). A permutation test for constrained multivariate analyses was performed to test the significance of discrimination between factors included in the RDA full model (experiment, isolate, subculturing host, and tested host). The full model was then simplified by excluding the factor “subculturing host”, as model comparison revealed no significant difference.

## Acknowledgments

The authors thank Mr. Robert Giovinazzo (Sonito, Avignon) for providing the 15.TII.02 isolate sample. They express their thanks to Christophe Piriou, Claudine Pasco and Bruno Marquer for technical assistance with the biological experiments, to the INRA Genetic Resources Center BrACySol of Ploudaniel for providing the potato seed tubers used in the tests, and to the IGEPP technical staff for managing greenhouse facilities. Useful discussions with and comments from Dr. Michel Ponchet (INRA ISA, Sophia Antipolis) are also gratefully acknowledged. This work was supported by the PoH-MED project, Potato Health – Managed for Efficiency and Durability, funded under the ARIMNet (Agricultural Research in the Mediterranean Area) ERA-Net (KBBE 219262).

## Supporting information captions

S1 Appendix. Hypothesis - The proteases inhibitors EPIC2B/EPI1 block immunity-related programmed cell death.

**S1 Fig. Effector genes equally expressed before and after experimental evolution.** Two *P. infestans* isolates collected on potato (P-isolate 15.P50.09) and on tomato (T-isolate 15.TII.02) had been subcultured nine times on their original host: the P-isolate on the potato cv. Bintje and the T-isolate on the tomato cv. Marmande. Effector gene expression was assessed before (dark grey bars) and after subculturing (white bars). Both measurements were performed on the potato cv. Bintje and on the tomato cv. Marmande, but results were averaged as the effects of subculturing were similar on both tested hosts. For effector genes shown here, pairwise t-tests revealed no significant differences in transcript abundance before and after nine times subculture (n=6, significance level α = 0.05, n.s. = not significant).

**S2 Fig. Loading plot of redundancy analysis.** Correlations among effector genes and fitness are displayed on the first and the third constrained component axis, representing respectively 56.54 and 15.48 % (cumulative 72.02 %) of the constraint variance.

**S3 Fig. Experimental evolution experiment design.** *Phytophthora infestans* inoculum was prepared on the host from which the isolate had been collected. This base inoculum served to start experimental evolution, consisting in nine times subculture on detached leaflets: either on the original host (from which the isolate had been collected), on the alternative host (e.g. on tomato leaflets if the isolate had been collected on potato), or on alternating hosts (in turns on potato and tomato). Each of these lineages was replicated three times. Fitness and effector gene expression of *P. infestans* on potato and tomato were assessed simultaneously, before and after experimental evolution.

S1 Table. *Phytophthora infestans* pathogenic fitness.

S2 Table. RT-PCRq primer sequences for genes of interest and reference genes, amplicon length and primer efficiency.

S3 Table. *Phytophthora infestans* effector gene expression (normalized transcript abundance).

S4 Table. *Phytophthora infestans* mean fitness and effector gene expression (normalized transcript abundance) for RDA analysis.

