## Supplemental Appendix 1 for "The level of specialization of *Phytophthora infestans* to potato and tomato is a biotrophy-related, stable trait"

**Hypothesis**

**The proteases inhibitors EPIC2B/EPI1 block immunity-related programmed cell death**

In the present study, the fitness of the pathogen *Phytophthora infestans* on potato and tomato was negatively related with the expression of the effectors EPIC2B and EPI1. Our discussion on the biological significance of this relationship – presented in the discussion section – is based on the hypothesis that *P. infestans* secretes EPIC2B and EPI1 to block immunity-related programmed cell death and regulate the host cell death machinery. To explain this hypothesis, we provide a simplified scheme (Fig 8. in the article) and hereafter a supporting review of literature.

The secretion of apoplastic protease inhibitors such as EPI1 and EPIC2B by *P. infestans* is understood as a counter-defense mechanism against the action of host proteases [1, 2], which are suspected to degrade pathogen effectors, pathogen structural molecules and/or to be involved in immune signalling [16, 17]. EPI1 is a Kazal-like serine protease inhibitor that inhibits the pathogenesis-related subtilisin-like apoplastic serine protease P69B (also known as PR-7) [2]; by inhibiting P69B, it therefore protects EPIC2B from degradation [3]. EPIC2B in turn inhibits the apoplastic tomato proteases C14 (also named SENU2/CYP1/TDI65) [4] and PIP1 (*Phytophthora* Inhibited Protease 1) [1]. Both are C1A Papain-Like Cysteine Proteinases (PLCPs, MEROPS database <http://merops.sanger.ac.uk>) and contribute to tomato immunity: C14 against *P. infestans* [4] + [5] and PIP1 against *Cladosporium fulvum* [6]. The precise role of C14 and PIP1 in tomato immunity is not understood, but could be related to their implication in Programmed Cell Death (PCD). PIP1 is indeed required for the Hypersensitive Cell Death (HCD) conferred by the tomato resistance gene *Cf-4* during the interaction with *Cladosporium fulvum* expressing the avirulence gene *Avr4* [6], while C14 is expressed in PCD events during seed germination and leaf senescence [7], as well as in tomato fruit stressed by extreme temperatures [8]. The tomato C14 protease has also been reported to be inhibited by the *Pseudomonas syringae* proteases inhibitor RIP1, that also inhibits the *Arabidopsis* vacuolar papain-like cysteine protease RD21 (Response to Desiccation 21) [4, see discussion section]. Interestingly, RD21 is the tomato C14 ortholog protease of *Arabidopsis*, and triggers cell death in this host [9]: RD21 knockout reduces PCD, while over expression enhances PCD. It is thus tempting to speculate that RIP1 from *Pseudomonas syringae* inhibits C14 in tomato – just as its ortholog RD21 in *Arabidopsis* – to block PCD.

Altogether, these literature findings strongly suggest that *P. infestans* secretes the proteases inhibitors EPI1 and EPIC2B to take control on the tomato cell death machinery. This may also be the case in the *P. infestans* – potato interaction, as a potato cysteine protease – highly homologous to tomato C14 – has been shown to accumulate more in highly resistant potato cultivars than in the susceptible cultivar Bintje [10].
