## Supplementary figures and images for "The level of specialization of *Phytophthora infestans* to potato and tomato is a biotrophy-related, stable trait"

### Supplemental Figure 1

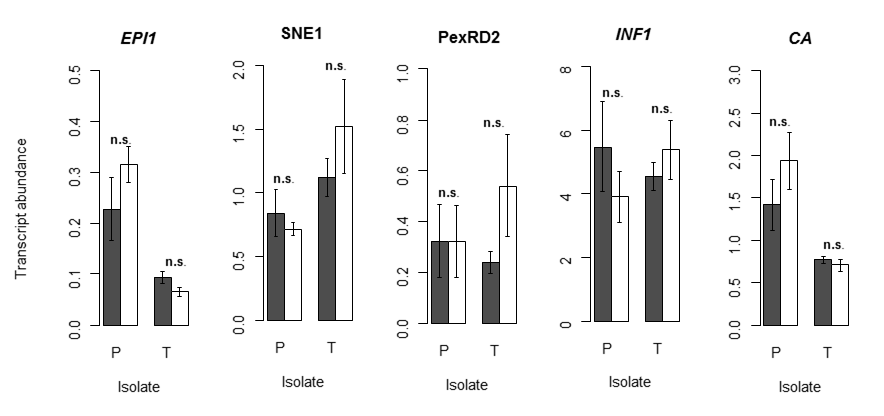

### Supplemental Figure 2

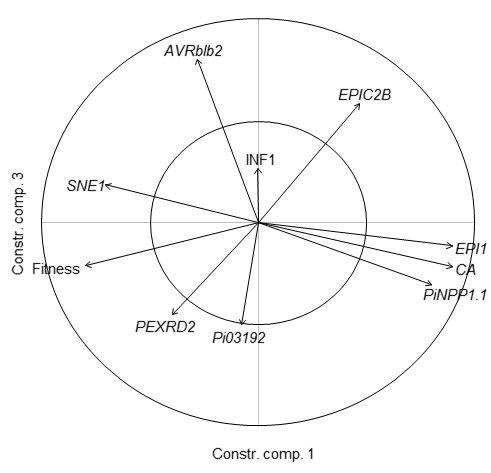

### Supplemental Figure 3

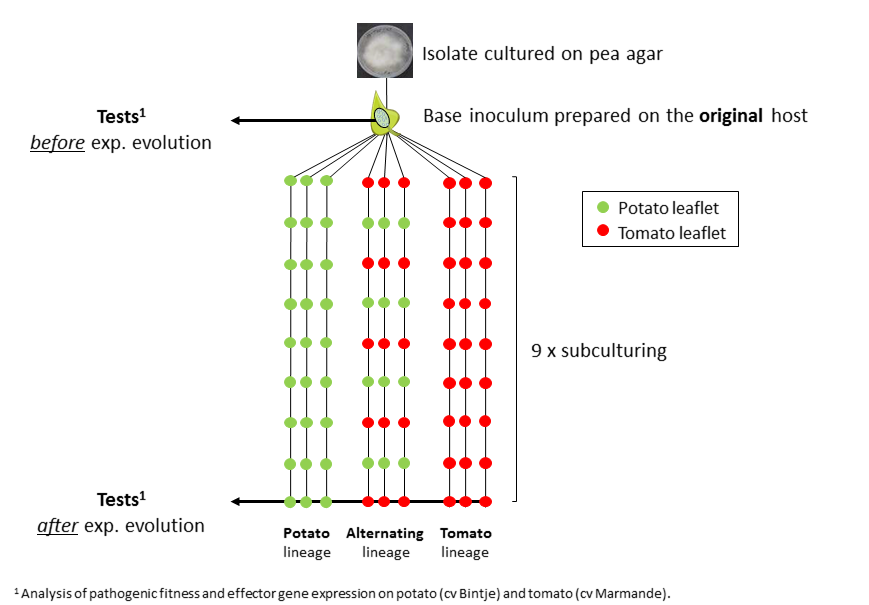
