## Supplemental Table 2 for "The level of specialization of *Phytophthora infestans* to potato and tomato is a biotrophy-related, stable trait"

**S1 Table: RT-qPCR primer sequences for genes of interest and reference genes, amplicon length and primer efficiency.**

| **Gene** | **PITG ID** | **Accession** | **Forward primer 5'-3'** | **Reverse primer 5'-3'** | **Amplicon (bp)** | **Efficiency ^1^ (%)** |
| --- | --- | --- | --- | --- | --- | --- |
| *AVRblb2* | PITG_04090 | XM_002905755 | GGATCGAGGCCCAAGAAGTT | CGGCCTCTTGACGATCTTGT | 101 | 83,6 |
| *EPIC2B* | PITG_09173 | XM_002903436 | GCCCAACTGAACGGATACTCA | GAGTTGACGCTGCAACCG | 197 | 82,7 |
| *EPI1* | PITG_22681 | XM_002907996 | CAAAGCCCGCAAGTCATCAG | CACTTTGCGCGCTTCATGTA | 146 | 81,2 |
| *PexRD2* | PITG_11384 | XM_002900934 | ATTGCGGCTTCATTCCTGGT | TCAGCTGCCGTGTAGTGTTT | 116 | 84,3 |
| *SNE1* | PITG_13157 | XM_002900631 | CTTGACAACCAGCAACGTCG | ATCGTTTTGGCTGGTGTTGC | 148 | 88,4 |
| *PiNPP1.1* | PITG_16866 | XM_002897255 | GTTCAAGCCCCAGATCCACA | TATCCGGAACCTTTGCACCC | 126 | 80,2 |
| *INF1* | PITG_12551 | XM_002900382 | CTTTCGTGCTCTGTTCGCTG | GAGTCCGTCGAGCACTGATT | 147 | 81,0 |
| *Pi03192* | PITG_03192 | XM_002906231 | AGCGGTGAAGAGAGAGCCTA | TGATGCCAGGCCTGCTAAAA | 67 | 80,6 |
| *Carbonic anhydrase* | PITG_14412 | XM_002898834 | AACCGACGAGGAATTGGGTC | TGTTGATGGGAGACTGGTGC | 80 | 81,6 |
| *Actin* | PITG_15117 | XM_002898238 | GACGTTCCAGCAGATGTGGA | CGTGGACAGGCAGCTTAGAA | 93 | 86,4 |
| *β-tubuline* | PITG_00156 | XM_002908737 | GGTCGTGGAGCCCTATAACG | GTGGTGAGCTTCAATGTGCG | 123 | 89,3 |
| *Elongation factor 2* | PITG_10941 | XM_002901697 | CCTCGTCACGACTTCAAGCT | GCCGTAGCCCCAGATCTTAC | 90 | 80,6 |

^1^ Mean PCRq efficiency (number of standard curves = 4).
